# Discovery of Alstrom syndrome gene as a regulator of centrosome duplication in asymmetrically dividing stem cells in *Drosophila*

**DOI:** 10.1101/490615

**Authors:** Cuie Chen, Yukiko M. Yamashita

## Abstract

Stereotypical inheritance of the mother vs. daughter centrosomes has been reported in several stem cells that divide asymmetrically. We report the identification of a protein that exhibits asymmetric localization between mother and daughter centrosomes in asymmetrically dividing *Drosophila* male germline stem cells (GSCs). We show that Alms1a, a *Drosophila* homolog of the causative gene for the human ciliopathy Alstrom Syndrome, is a ubiquitous mother centriole protein with a unique additional localization to the daughter centriole only in the mother centrosome of GSCs. Depletion of *alms1a* results in rapid loss of centrosomes due to failure in daughter centriole duplication. We reveal that *alms1a* is specifically required for centriole duplication in asymmetrically dividing cells but not in symmetrically dividing differentiating cells in multiple stem cell lineages. The unique requirement of *alms1a* in asymmetric dividing cells may shed light onto the molecular mechanisms of Alstrom syndrome pathogenesis.

## INTRODUCTION

Many stem cells divide asymmetrically to generate daughter cells with distinct fates: one self-renewing stem cell and one differentiating cell. Asymmetrically dividing stem cells in several systems show a stereotypical inheritance of the mother vs. daughter centrosomes (Conduit and Raff, 2010; Habib et al., 2013; Januschke et al., 2011; Wang et al., 2009; Yamashita et al., 2007). This observation has provoked ideas that stem cell centrosomes might play a critical role in achieving cell fate asymmetry and might be uniquely regulated. The first step toward testing these ideas is to discover mother or daughter centrosome-specific proteins that exhibit stem cell-specific functionality in asymmetrically dividing stem cells, yet no such proteins have been identified to date.

*Drosophila* male germline stem cells (GSCs) divide asymmetrically by orienting the spindle perpendicular toward the hub cells, the major niche component (Yamashita et al., 2003) (Figure 1A). The stereotypical behavior of mother vs. daughter centrosomes was first discovered in the *Drosophila* male GSC (Yamashita et al., 2007): the mother centrosome is always located near the hub-GSC junction, whereas the daughter centrosome migrates to the other side of the cell, leading to spindle orientation perpendicular to the hub and consistent inheritance of the mother centrosome by GSCs (Yamashita et al., 2007).

**Figure 1.**
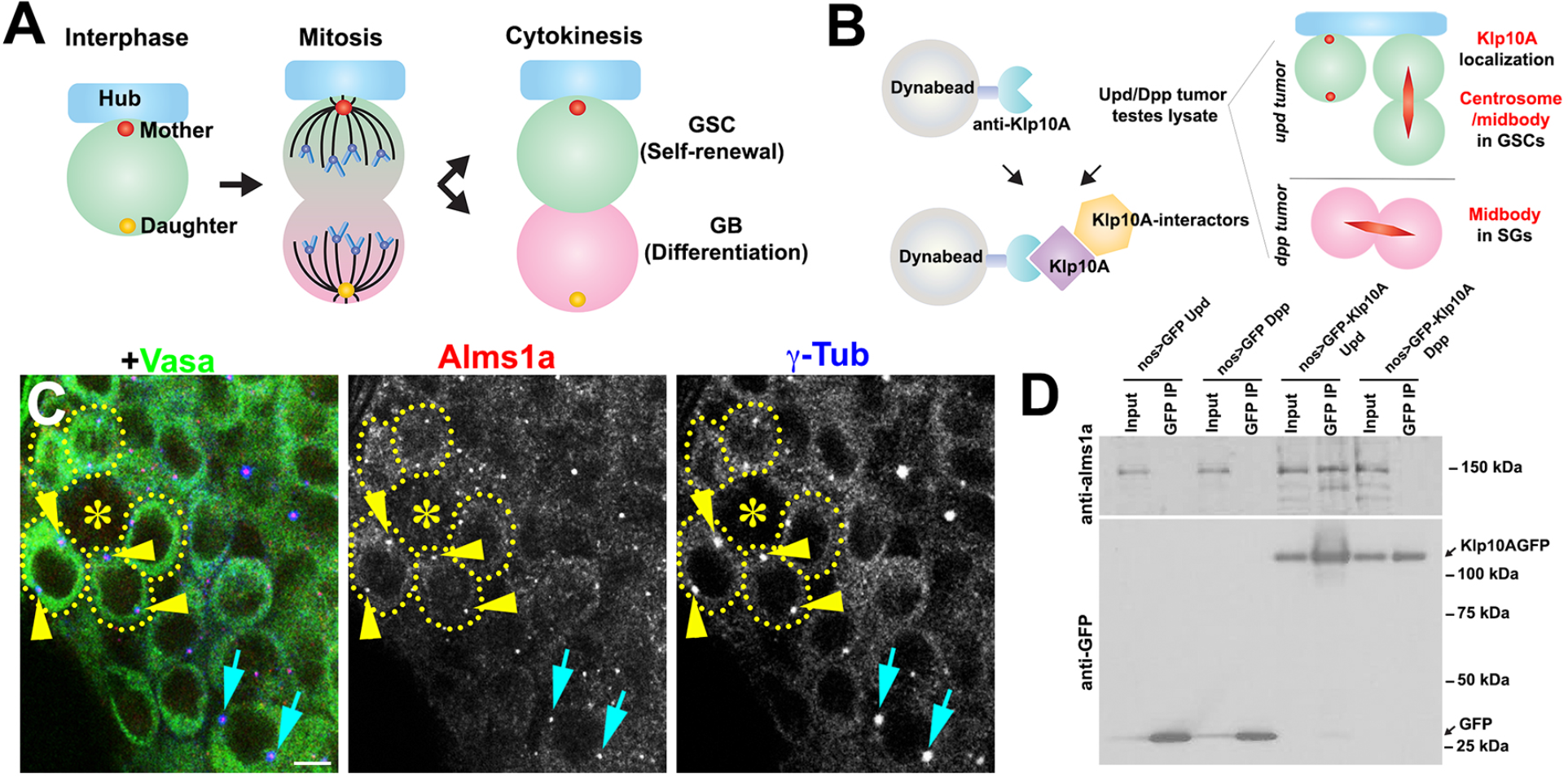
Identification of Alms1a as a germline-stem cell-specific Klp10A interactor. (A) Asymmetric centrosome inheritance in Drosophila male GSC. (B) Scheme of Klp10A pulldown and mass spectrometry. Klp10A pull down was conducted using either *upd*-induced tumor (GSC-enriched) or *dpp*-induced tumor (SG-enriched), followed by mass spectrometry analysis. (C) An apical tip of the *Drosophila* testis stained for Alms1a (red), Blue: γ-Tub (centrosome/pericentriolar matrix) and Green: Vasa (germ cells). Asterisk indicates the hub. GSCs are outlined with yellow dotted lines. Arrowheads (yellow) indicate examples of GSC centrosomes. Arrows (cyan) indicate examples of SG centrosomes. Bar: 5 μm. (D) Co-immunoprecipitation of Klp10A and Alms1a. Control GFP and GFP-Klp10A was pulled down from GSC extracts using an anti-GFP antibody and blotted with anti-Alms1a and anti-GFP.

To obtain insights into the underlying mechanism/regulation of asymmetric centrosome inheritance, we attempted to identify proteins that exhibit mother or daughter-centrosome specific localization in GSCs. Here, we identify Alms1a, a *Drosophila* homolog of the causative gene for the human ciliopathy Alstrom Syndrome, as a centrosomal protein that exhibit unique asymmetric localization between mother and daughter centrosomes in GSCs. *alms1a* depletion leads to failure in daughter centriole duplication in GSCs, resulting in loss of centrosomes in GSCs as well as all differentiating daughters of GSCs. Strikingly, *alms1a*’s requirement in centriole duplication is specific to GSCs, and symmetrically dividing differentiating cells or GSCs induced to divide symmetrically do not require *alms1a* for centriole duplication. We further show that two other asymmetrically dividing stem cells in *Drosophila*, neuroblasts and female GSCs exhibit similar *alms1a*-dependence for centriole duplication. Our study may provide a link between asymmetric stem cell division and the pathogenesis of Alstrom Syndrome.

## RESULTS

### Identification of Alms1a as a GSC-specific Klp10A interactor

Previously, we reported that Klp10A, a microtubule-depolymerizing kinesin of kinesin-13 family, is enriched on the centrosomes of GSCs, but not on the centrosomes of differentiating germ cells (e.g. gonialblasts (GBs) and spermatogonia (SGs)) (Chen et al., 2016). Depletion of Klp10A in GSCs results in overly elongated mother centrosomes and slightly smaller daughter centrosomes in GSCs but does not affect centrosomes in differentiating GBs/SGs, suggesting a specialized regulation of centrosomes in GSCs (Chen et al., 2016). We reasoned that identifying Klp10A-interacting proteins may uncover proteins that regulate the asymmetric behavior of mother and/or daughter centrosomes in GSCs. In addition to the GSC centrosomes, Klp10A localizes to the central spindle of all germ cells (Chen et al., 2016). To enrich for Klp10A-interactors on GSC centrosomes but not on central spindles (Figure 1B), we conducted parallel immunopurification-mass spectrometry analysis comparing GSC extract vs. SG extract (Figure 1B). We immunopurified Klp10A from testis extract enriched for GSCs due to overexpression of the self-renewal factor, Upd, a cytokine-like ligand (Kiger et al., 2001; Tulina and Matunis, 2001) and from testis extract enriched for SGs due to overexpression of Dpp, a bone morphogenic protein (BMP) (Bunt and Hime, 2004; Kawase et al., 2004; Schulz et al., 2004). We subtracted proteins that co-immunopurified with Klp10A between GSC-enriched testes and SG-enriched testes to enrich for those that localize specifically to GSC centrosomes.

Using this approach, we identified 260 candidates (Supplementary data file). Of these, *Alstrom syndrome (Alms) 1* stood out to us, as its mammalian homolog was previously shown to localize to centrosomes and basal bodies of the primary cilia (Andersen et al., 2003; Hearn et al., 2005). Alstrom syndrome is a rare autosomal recessive disorder characterized by childhood obesity and sensory impairment, and categorized as multiorgan ciliopathy (Marshall et al., 2011). The *Drosophila* genome encodes two *Alms1* homologs, juxtaposed next to each other on the X chromosome, which we named *alms1a (CG12179*) and *alms1b (CG12184)*. The peptide sequences identified by our mass-spectrometry analysis were identical between these two proteins, so to discern whether one or both homologs interact with Klp10A in GSCs, we raised specific antibodies against Alms1a and Alms1b. We found that Alms1a localized to centrosomes broadly in early germ cells (GSCs to spermatocytes) (Figures 1C and S1A), whereas Alms1b was not expressed in these early germ cells but localized to the basal body of elongating spermatids (Figures S1C and S1D). Therefore, Alms1a is likely the relevant Klp10A interactor in GSCs. Moreover, Alms1a and Klp10A co-immunoprecipitated in GSC-enriched extract but not in SG-enriched extract, demonstrating that Alms1a is a GSC-specific Klp10A interactor (Figure 1D). Therefore, although Alms1a localizes to the centrosomes of both GSCs and SGs, Alms1a might have a specific function in GSCs via its interaction with Klp10A.

### Alms1a localizes asymmetrically between mother versus daughter centrosome in male GSCs

To understand the function of Alms1a, possibly specific to GSCs, we first examined its centrosomal localization in GSCs and SGs in more detail. Each centrosome consists of two centrioles, the mother centriole (‘m’ in Figure 2A) and daughter centriole (‘d’ in Figure 2A). The centrosome that contains the older mother centriole is called the mother centrosome (‘M’ in Figure 2A), whereas the centrosome that contains the younger mother centriole is called daughter centrosome (‘D’ in Figure 2A).

**Figure 2.**
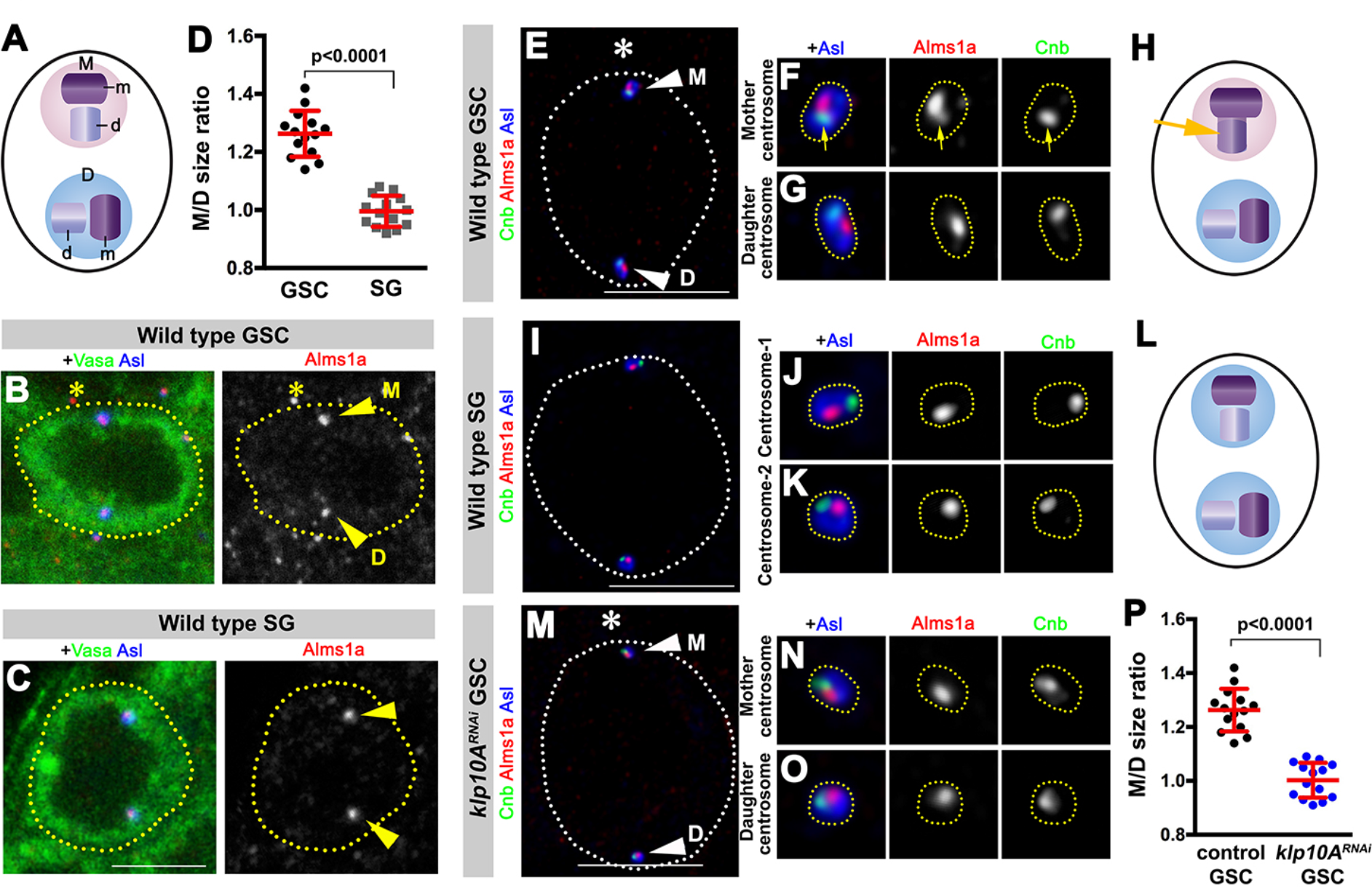
Alms1a is a mother centriole protein with additional localization to the daughter centriole only on GSC mother centrosomes. (A) Each GSC contains two centrosomes, mother centrosome (M, pink) and daughter centrosome (D, blue). Each centrosome consists of two centrioles, mother centriole (m, dark purple) and daughter centriole (d, light purple). (B and C) Alms1a localization in wild type GSC (B) and SG (C). Green: Vasa. Red: Alms1a. Blue: Asl (centriole). Asterisk indicates the hub. Arrowheads indicate centrosomes. Bar: 5 μm. (D) Quantification of Alms1a size ratio (Mother/Daughter centrosome, M/D) in the indicated cell types. P value was calculated using two-tailed Student’s t-test. Error bars indicate the standard deviation. (E-H) A HyVolution image of wild type GSC centrosomes (E). (F and G) Magnified images of a mother centrosome (F) and a daughter centrosome (G) from panel (E). Green: Centrobin (Cnb). Red: Alms1a. Blue: Asl. Asterisk indicates the hub. Arrowheads indicate GSC centrosomes. Arrow indicates mother centrosome. Bar: 5 μm. (H) A graphic interpretation. Arrow indicates Alms1a’s localization to the daughter centriole of the GSC mother centrosome. (I-L) A hyvolutoin image of wild type SG centrosomes (I). (J and K) Magnified images of SG centrosomes from panel (I). Green: Cnb. Red: Alms1a. Blue: Asl. Asterisk indicates the hub. Bar: 5 μm. (L), A graphic interpretation. (M-O) A HyVolution image of *klp10A*^*RNAi*^ GSC centrosomes (M). (N and O) Magnified images of a mother centrosome (N) and a daughter centrosome (O) from panel (M). Green: Centrobin (Cnb). Red: Alms1a. Blue: Asl. Asterisk indicates the hub. Arrowheads indicate GSC centrosomes. Bar: 5 μm. (P) Quantification of Alms1a size ratio between mother and daughter centrosomes (M/D) of GSCs in the indicated genotypes. P value was calculated using two-tailed Student’s t-test. Error bars indicate the standard deviation

Importantly, we found that Alms1a exhibited an asymmetric localization to centrosomes specifically in GSCs. In GSCs, the mother centrosome, which is located near the hub cells, consistently had more Alms1a compared to the daughter centrosome (Figures 2B and 2D). In contrast, the two centrosomes in SGs had an equivalent amount of Alms1a (Figures 2C and 2D). Detailed examination of Alms1a localization using HyVolution confocal microscopy (see Methods) revealed that Alms1a is a mother centriole protein in GSCs and SGs, exhibiting non-overlapping localization with the daughter centriole protein Centrobin (Cnb) (Januschke et al., 2013) (Figures 2E-2L). Interestingly, Alms1a additionally localized to the daughter centriole specifically at the mother centrosome in GSCs (Figures 2E, 2F and 2H), accounting for the higher amount of Alms1a on GSC mother centrosomes (Figures 2B and 2D).

To determine if *klp10A* is required for asymmetric localization of Alms1a in GSCs, we depleted *klp10A* in the germline by RNAi. Knockdown of *klp10A* in male germline (*nos-gal4>klp10A*^*TRiP.HMS00920*^, *klp10A*^*RNAi*^ hereafter (Chen et al., 2016)) abolished Alms1a localization to the daughter centriole of the mother centrosome in GSCs (Figures 2M-2O), leaving an equal amount of Alms1a on the mother centrioles of both centrosomes in GSCs (Figure 2P). *klp10A*^*RNAi*^ did not affect Alms1a localization to the SG centrosomes (Figure S2C). Germline knockdown of *alms1a* (*nos-gal4>alms1*^*TRiP.HMJ30289*^, referred as to *alms1*^*RNAi*^ hereafter) did not compromise Klp10A localization in GSCs (Figures S2A and S2B). These results, combined with the GSC-specific physical interaction with Klp10A (Figure 1D), show that the Klp10A-Alms1a interaction mediates the recruitment of Alms1a to the daughter centriole of the GSC mother centrosome and indicate that relevant interaction of Klp10A-Alms1a occurs on the daughter centriole of the GSC mother centrosome.

### Multiple types of *Drosophila* stem cells exhibit asymmetric Alms1a localization between the mother and daughter centrosomes

To examine whether Alms1a exhibit asymmetric localization between the mother and daughter centrosomes in other asymmetrically dividing stem cells, we examined 3rd instar larval neuroblast and female GSCs (Figure 3). Both in neuroblasts and female GSCs, the mother centrosomes had more Alms1a compared to the daughter centrosome, due to the presence of Alms1a to the daughter centrioles of the mother centrosomes (Figures 3A-3C, 3G, 3H-3J and 3N). On the contrary, differentiating cells of their lineage (cystocytes and ganglion mother cells (GMCs)) did not show any asymmetry in Alms1a amount between two centrosomes (Figures 3D-3G, and 3K-3N). It is interesting to note that both female GSCs and neuroblasts inherit the daughter centrosomes, whereas the mother centrosomes are segregated to differentiating cells (Januschke et al., 2011; Salzmann et al., 2014). In the neuroblasts, during the asymmetric inheritance of the mother vs. daughter centrosomes, the mother centrosome is functionally inactivated as a microtubule-organizing center (MTOC) until mitotic entry, whereas the daughter centrosome retains robust MTOC activity (Conduit and Raff, 2010; Rebollo et al., 2007; Rusan and Peifer, 2007). Therefore, Alms1a’s localization to the daughter centriole of the mother centrosome is unlikely correlated with their MTOC activity. Taken together, these results indicate that the asymmetric centrosomal localization of Alms1a extends to diverse types of asymmetrically dividing stem cells.

**Figure 3.**
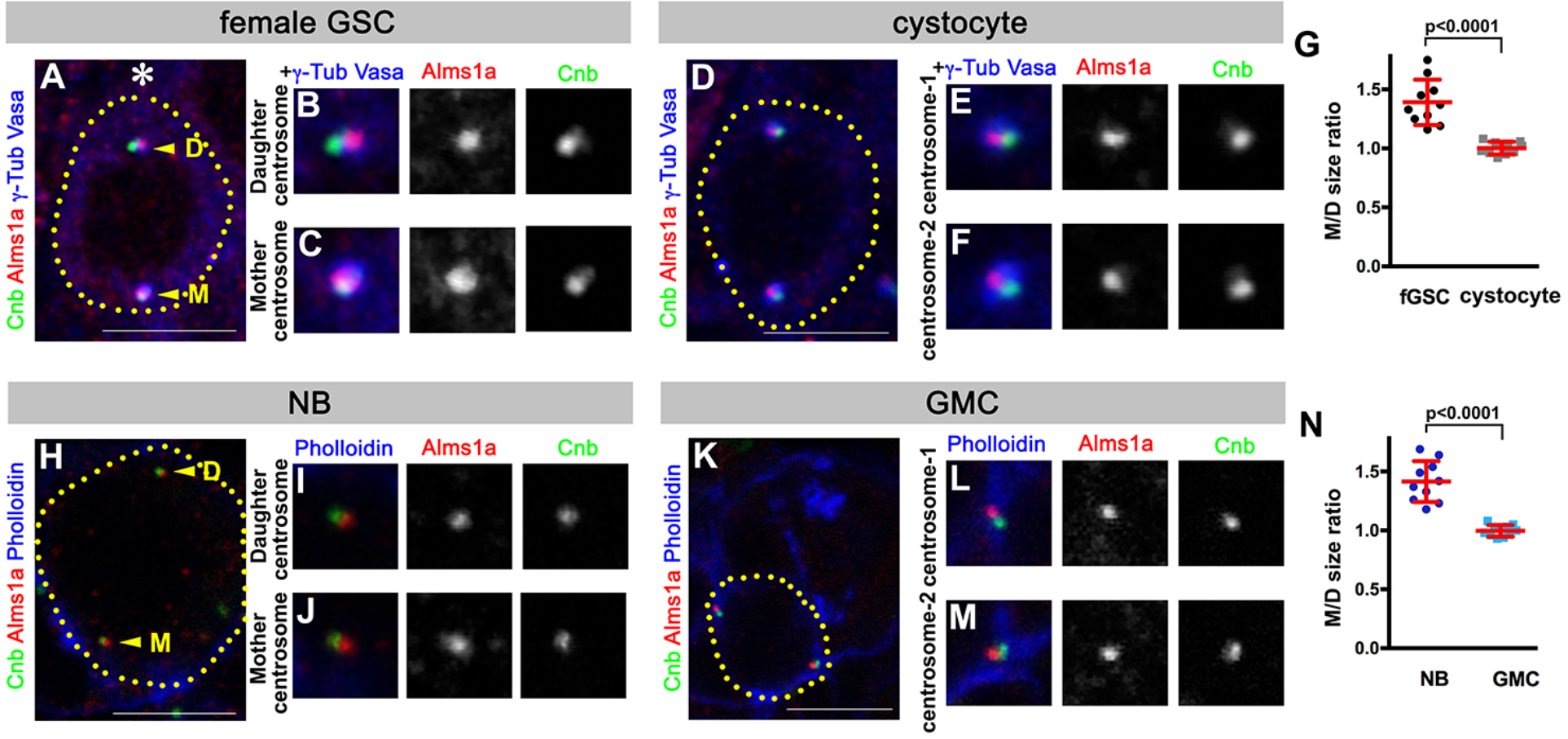
Alms1a localization in female GSC and neuroblast (NB) (A-C) An example of wild type female GSC centrosomes (A). (B and C) Magnified images of a daughter centrosome (B) and a mother centrosome (C) from panel (A). Green: Cnb. Red: Alms1a. Blue: γ-Tub and Vasa. Asterisk indicates the niche. Arrowheads indicate centrosomes. Bar: 5 μm. (D-F) An example of wild type cystocyte (differentiating germ cell) centrosomes (D). (E and F) Magnified images of two centrosomes from panel (D). Green: Cnb. Red: Alms1a. Blue: γ-Tub and Vasa. Bar: 5 μm. (G) Quantification of Alms1a size ratio (Mother/Daughter centrosome, M/D) in the female GSCs abd cystocytes. P value was calculated using two-tailed Student’s t-test. Error bars indicate the standard deviation. (H-J) An example of wild type NB centrosomes (H). (I-J) Magnified images of a daughter centrosome (I) and a mother centrosome (J) from panel (H). Green: Cnb. Red: Alms1a. Blue: Phalloidin. Arrowheads indicate centrosomes. Bar: 5 μm. (K-M) An example of wild type GMC (ganglion mother cell, differentiating daughter of NB) centrosomes (K). (L and M) Magnified images of two centrosomes from panel (K). Green: Cnb. Red: Alms1a. Blue: Phalloidin. Bar: 5 μm. (N) Quantification of Alms1a size ratio (Mother/Daughter centrosome, M/D) in the NBs and GMCs. P value was calculated using two-tailed Student’s t-test. Error bars indicate the standard deviation.

### Depletion of *alms1a* leads to centrosome loss due to defective centriole duplication

To understand the function of *alms1a*, we examined the phenotypes upon depletion of *alms1a*. Depletion of *alms1a* was confirmed using a specific antibody against Alms1a (Figures S1A, S1B and S1F). Note that *UAS-alms1*^*RNAi*^ is expected to knockdown both *alms1a* and *alms1b*. However, because *alms1b* is not expressed in GSCs/SGs (Figure S1C), the phenotypes caused by *alms1*^*RNAi*^ in GSCs are most likely due to the loss of *alms1a* (see Methods).

Knockdown of *alms1a* resulted in a rapid loss of centrosomes from all germ cells, except for the mother centrosomes in GSCs, demonstrating that *alms1a* is required for centrosome duplication but not the maintenance of existing centrosomes (Figures 4A and 4B). Wild type GSCs normally contain both mother and daughter centrosomes throughout the cell cycle, due to early duplication of the centrosome right after the previous mitosis (Figure 4C). In contrast, most *alms1*^*RNAi*^ GSCs contained only one centrosome, visualized by Asl as well as γ-Tubulin (Figures 4E and 4F). Both wild type and *alms1*^*RNAi*^ GSCs underwent bipolar mitotic divisions: in *alms1*^*RNAi*^ GSCs with one centrosome remaining, the centrosome was consistently observed at the proximal side of mitotic spindle, whereas the other side lacked the centrosome (Figures 4D and 4F). Interestingly, the mother centrosome in GSCs gradually elongated upon knockdown of *alms1a* (Figures 4E, and S3), as we previously observed with *klp10A*^*RNAi*^ (Chen et al., 2016). These data suggest that *klp10A* and *alms1a* share a similar function, possibly balancing the growth of mother vs. daughter centrosomes. Taken together, we conclude that *alms1a* is required for centrosome duplication in GSCs.

**Figure 4.**
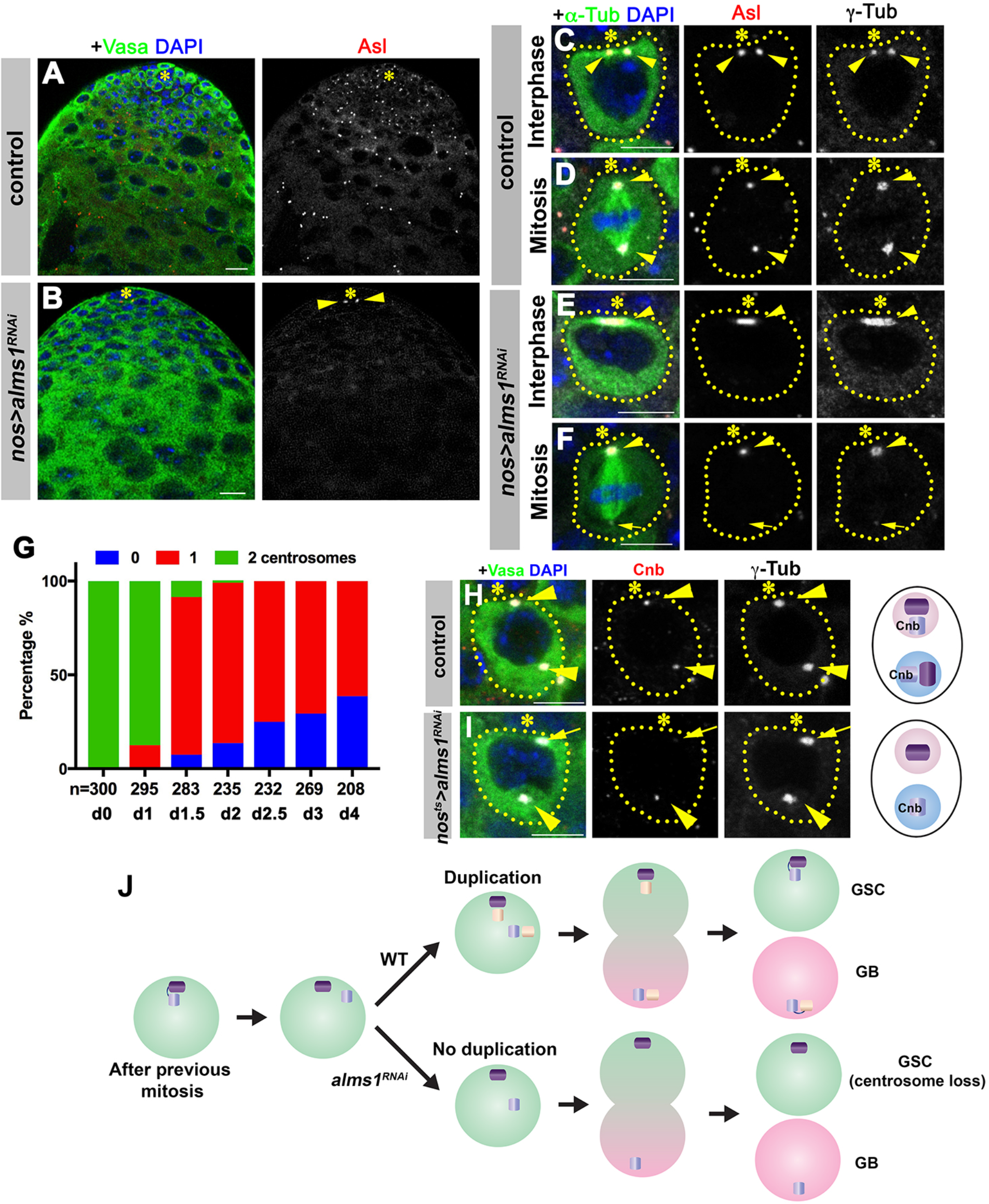
*alms1a* is required for daughter centriole duplication in GSCs. (A and B) Examples of apical tips in control (A) and *nos-gal4>UAS-alms1*^*RNAi*^ (B) testes. Green: Vasa. Red: Asl. Blue: DAPI. Asterisk indicates the hub. Arrowheads indicate remaining centrosomes in *alms1*^*RNAi*^ GSCs. Bar: 5 μm. (C-F) Examples of centrosomes in control interphase (C), control mitotic (D), *alms1*^*RNAi*^ interphase (E) and *alms1*^*RNAi*^ mitotic (F) GSCs. Green: GFP-α-tubulin. Red: Asl. White: γ-Tub. Blue: DAPI. Asterisk indicates the hub. GSCs are outlined with yellow dotted lines. Arrowheads indicate GSC centrosomes. Bar: 5 μm. (G) Quantification of centrosome number in temporarily controlled *alms1*^*RNAi*^ GSCs. (0-4 d after RNAi induction, *nos-gal4, tub-gal80*^*ts*^ >*UAS-alms1*^*RNAi*^). Centrosome number was quantified by Asl staining. (H and I) Examples of GSCs in control GSC (H) and *alms1*^*RNAi*^ GSC (I, 2 d after RNAi induction) stained for daughter centriole protein Cnb. Green: Vasa. Red: Cnb. White: γ-Tub. Blue: DAPI. Asterisk indicates the hub. Arrowheads indicate GSC centrosomes. Arrow indicates mother centrosome in *alms1*^*RNAi*^ GSC lacking Cnb staining (2d after RNAi induction). Cnb localization to the daughter centriole is indicated by ‘Cnb’ in the cartoon representations. Bar: 5 μm. N=30 GSCs for each genotype. (J) Model of centrosome loss in *alms1*^*RNAi*^ GSCs.

To understand the precise steps toward daughter centrosome loss upon depletion of *alms1a*, we induced germline *alms1*^*RNAi*^ in a temporarily controlled manner (*nos-gal4ΔVP16, tub-gal80*^*ts*^>*UAS-alms1*^*RNAi*^). This method confirmed that GSCs rapidly lose daughter centrosomes upon depletion of *alms1a* (Figure 4G), and allowed us to examine centrosomes prior to daughter centrosome loss. In control GSCs, the daughter centriole protein Cnb always marked both mother and daughter centrosomes, as expected because both centrosomes contain daughter centrioles (Figure 4H). However, upon knockdown of *alms1a*, we observed Cnb loss from the mother centrosome in GSCs, revealing that the GSC mother centrosome has lost the daughter centriole (Figure 4I). This indicates a failure of the mother centriole to produce a new (daughter) centriole in *alms1*^*RNAi*^ GSCs: upon loss of *alms1a*, the mother centriole (lacking Cnb) and daughter centriole (marked with Cnb) within the mother centrosome split from each other and segregate to GSC and GB, respectively, without having duplicated new centrioles, eventually leading to the centrosome loss in differentiating cells (Figures 4I and 4J). Without receiving the templating centrosomes from GSCs, differentiating germ cells cannot generate centrosomes, resulting in the complete loss of centrosomes/centrioles in all differentiating germ cells (Figure 4B). In the absence of centrioles, *nos-gal4>UAS-alms1*^*RNAi*^ flies were completely sterile (Figure S4). Note that the sterility of *nos-gal4>UAS-alms1*^*RNAi*^ flies likely reflects the depletion of both *alms1a* and *alms1b* throughout the spermatogenesis: however, the depletion of *alms1a* and *alms1b* only in differentiating germ cells (*bam-gal4>UAS-alms1*^*RNAi*^) leads to milder fertility defect, revealing the importance of alms1a in centriole duplication in GSCs. Taken together, we conclude that *alms1a* plays a critical role in ensuring centrosome duplication in GSCs, which is essential for production of centrosomes/centrioles in all downstream germ cells.

Similar to male GSCs, female GSCs and neuroblasts (NBs) also quickly lost centrosomes upon knockdown of *alms1a* (Figure 5). The centrosome loss upon depletion of *alms1a* in these cell types was much more profound compared to *alms1a*-depleted male GSCs (compare Figure 4G with Figures 5C and 5F). This is likely due to the fact that the mother centrosomes are inherited by differentiating cells in these cell types (Rebollo et al., 2007; Rusan and Peifer, 2007). Upon depletion of *alms1a*, daughter centrosomes fail to duplicate and the remaining mother centrosomes are inherited by differentiating cells, leaving no centrosomes in these stem cells. Together, we conclude that *alms1a* is required for centriole duplication in diverse stem cells.

**Figure 5.**
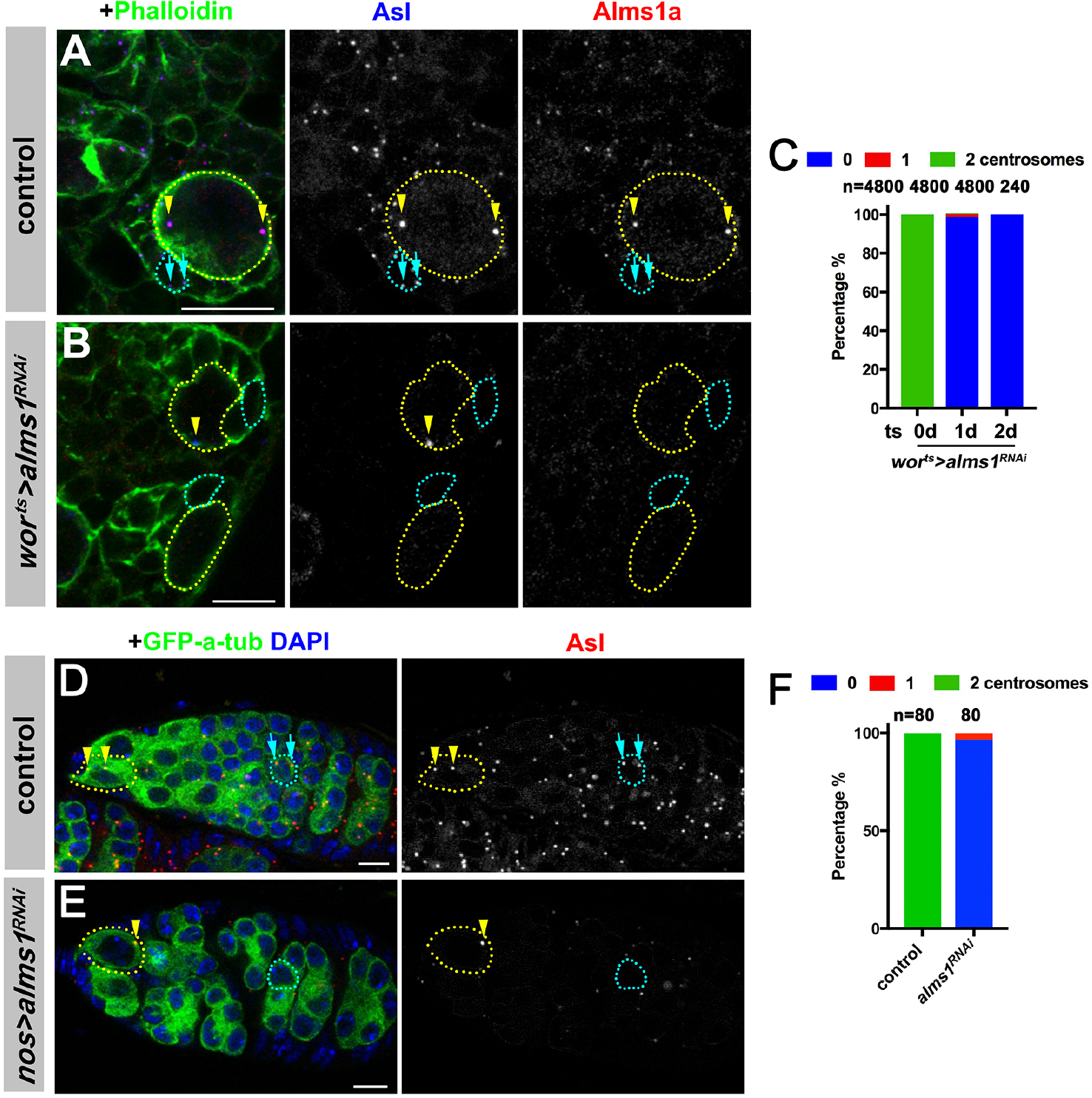
*alms1a* is required for centrosome duplication in female GSCs and NBs. (A and B) Examples of NBs in control (A) and *wor-gal4, gal80*^*ts*^>*UAS-alms1*^*RNAi*^ (1 d after RNAi induction) (B) brains. Green: Phalloidin. Red: Alms1a. Blue: Asl. Yellow dotted lines indicate NBs. Arrowheads indicate NB centrosomes. Cyan dotted lines indicate examples of GMCs. Arrows indicate GMC centrosomes. Bar: 10 μm. (C) Quantification of centrosome numbers in NBs in the indicated genotypes. (D and E) Examples of control (D) and *nos-gal4>UAS-alms1*^*RNAi*^ (E) germaria. Green: GFP-α-tubulin. Yellow dotted lines indicate examples of GSCs. Arrowheads indicate GSC centrosomes. Cyan dotted lines indicate examples of cystocysts. Arrows indicate cystocyst centrosomes. Red: Asl. Blue: DAPI. Bar: 5 μm. (F) Quantification of centrosome numbers in female GSCs of the indicated genotypes.

### *alms1a* is dispensable for centrosome duplication in symmetrically dividing, differentiating cells

The above results show that *alms1a* is required to ensure the duplication of daughter centrioles in multiple cell types that are known to divide asymmetrically. Considering the unique asymmetric localization of Alms1a only in stem cells (male and female GSCs and NBs) but not in differentiating cells, we wondered whether Alms1a is generally required for centriole duplication in differentiating cells.

To test this possibility, we depleted *alms1a* from differentiating cells in the testis as well as in the brain. SG-specific driver *bam-gal4* was used to deplete *alms1a i*n differentiating germ cells (*bam-gal4>UAS-alms1*^*RNAi*^). Similarly, *erm-gal4* was used to knockdown *alms1a* specifically in differentiating daughter of NBs, i.e. intermediate neural progenitors (INPs) (*erm-gal4>UAS-GFP, UAS-alms1*^*RNAi*^). There are two classes of NBs, type I, which produces GMCs upon division, and type II, which produces INPs. Since GMC-specific driver is not available, we focused on INP, where specific driver, *erm-gal4*, is available. Note that although the divisions of INPs are asymmetric in fate, generating another INP and one GMCs, their division apparatus (spindle) is symmetric (Bello et al., 2008; Boone and Doe, 2008; Bowman et al., 2008).

We found that, when *alms1a* was depleted from symmetrically dividing SGs, the SGs maintained their centrosomes, although Alms1a protein was efficiently depleted (Figures 6A and 6B). Similarly, knockdown of *alms1a* in INPs did not result in centrosome loss despite efficient depletion (Figures 6C and 6D). These results demonstrate that *alms1a* is specifically required for centriole duplication in asymmetrically dividing stem cells, but not in symmetrically dividing, differentiating cells.

**Figure 6.**
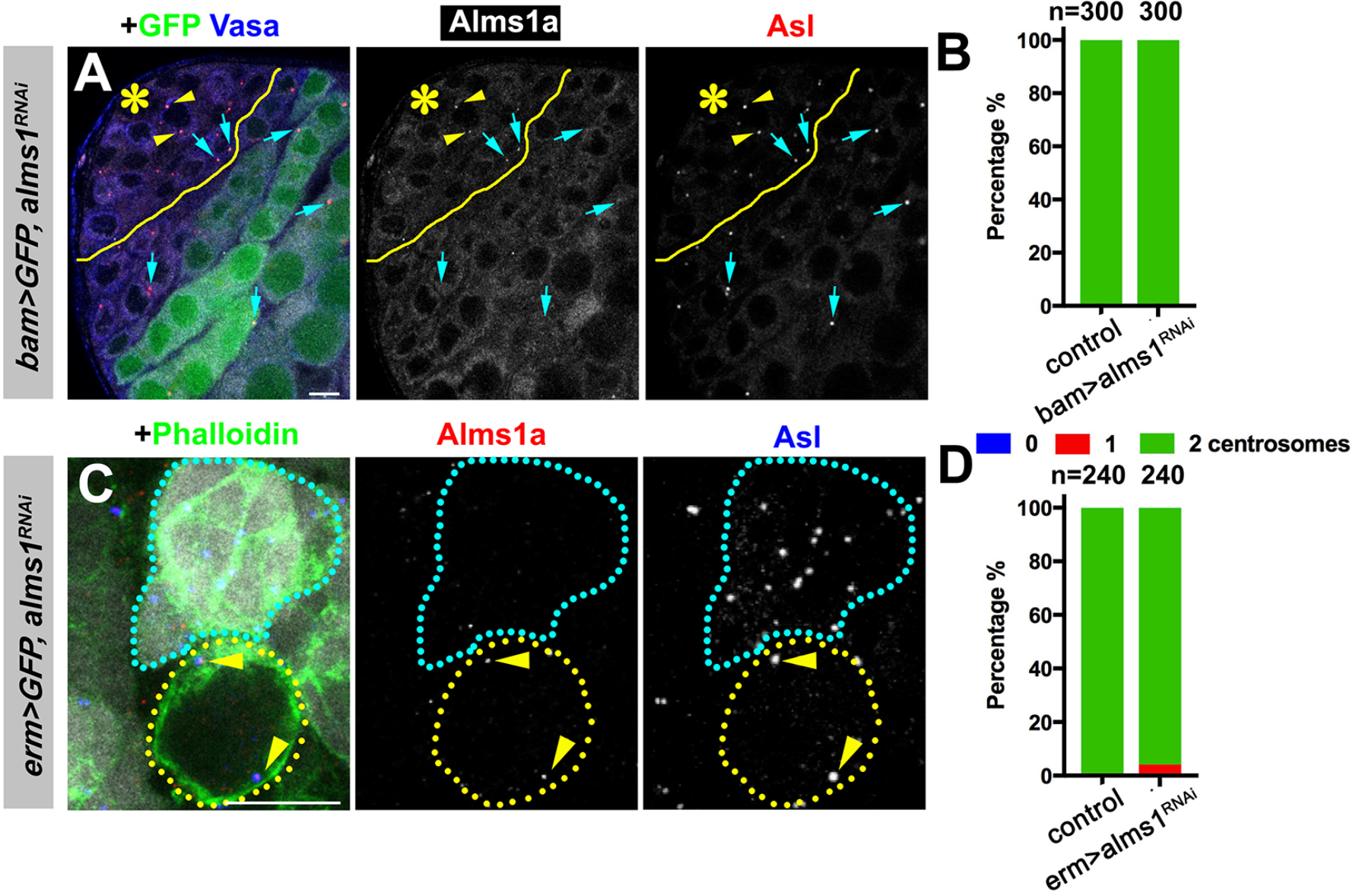
*alms1a* is dispensable for centrosome duplication in symmetrically dividing, differentiating cells. (A) An example of Alms1a staining in *bam-gal4>UAS-GFP, UAS-alms1*^*RNAi*^ testis. The boundary of bam-gal4-positive vs. −negative SGs is indicated by the yellow solid line. Green: GFP. White: Alms1a. Red: Asl. Blue: Vasa. Asterisk indicates the hub. Arrowheads indicate examples of GSC centrosomes. Arrow indicates examples of SG centrosomes. Bar: 10 μm. (B) Quantification of centrosome numbers in the indicated genotypes. (C) An example of Alms1a staining in *erm-gal4>UAS-GFP, UAS-alms1*^*RNAi*^ brains. Green: Phalloidin. White: GFP. Red: Alms1a. Blue: Asl. NBs are outlined with yellow dotted lines. Arrowheads indicate NB centrosomes. Cyan dotted line indicates NB daughters, in which Alms1a is depleted. Despite efficient depletion of Alms1a, these cells retain centrosomes (Asl). Bar: 10 μm. (D) Quantification of centrosome numbers in the indicated genotypes.

### *alms1a* is dispensable for centriole duplication in stem cells that are induced to divide symmetrically

The above results indicate that *alms1a* is required for centriole duplication uniquely in asymmetrically dividing stem cells, because *alms1a* depletion in symmetrically dividing, differentiating cells did not result in centriole duplication defect. To address whether the requirement of *alms1a* is linked to asymmetric division or stem cell identity, we examined whether symmetrically dividing stem cells require *alms1a* for centriole duplication (Figures 7A and 7B). We induced symmetric divisions to male GSCs by expressing *Upd* together with *alms1*^*RNAi*^ (*nos-gal4>UAS-alms1*^*RNAi*^, *UAS-Upd*). Alms1a protein was undetectable under this condition, confirming efficient knockdown (Figure 7B), yet these symmetrically-dividing GSCs retained centrosomes (Figure 7C), suggesting that these symmetrically-dividing GSCs do not require *alms1a* for centriole duplication.

**Figure 7.**
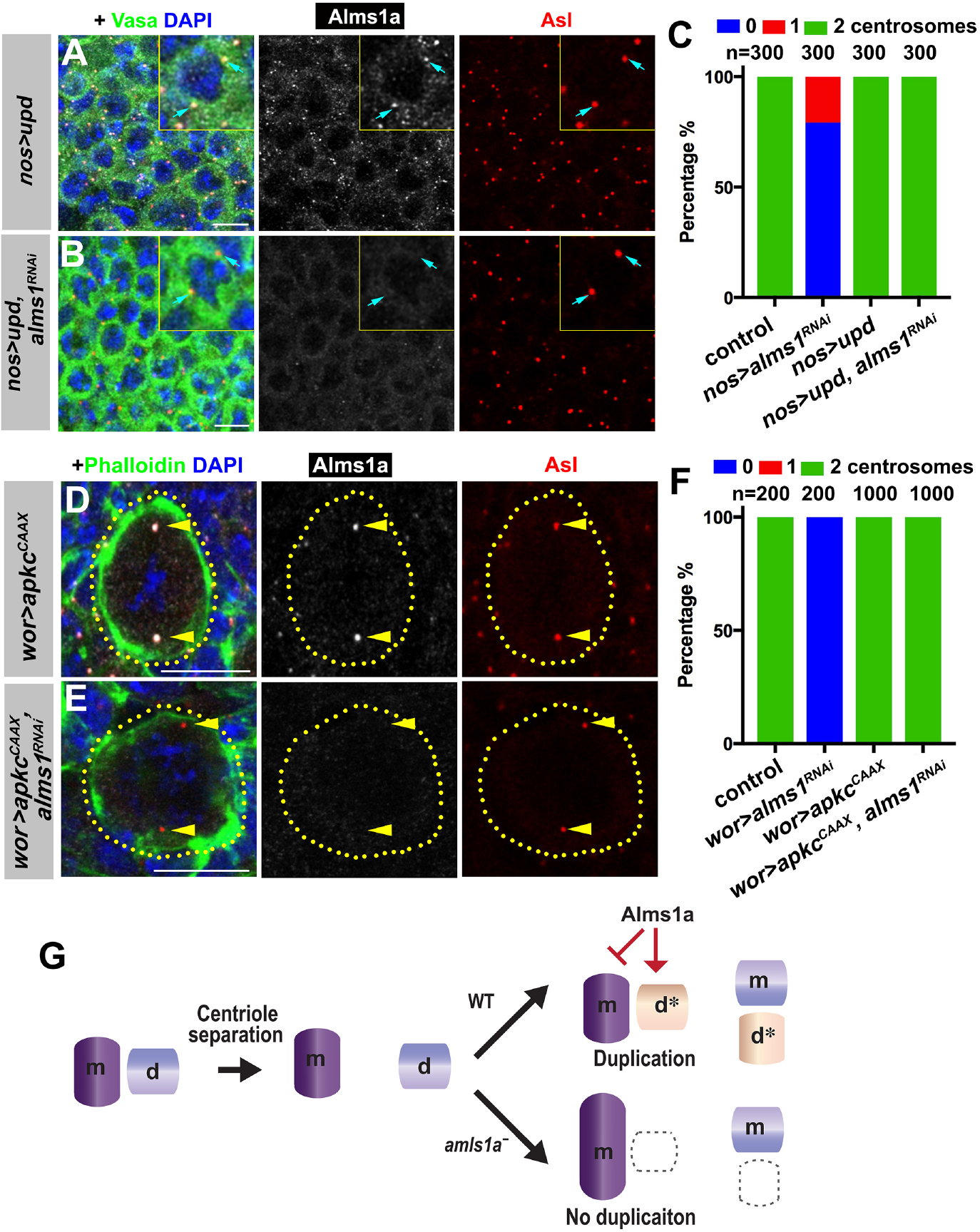
*alms1a* is not required for centrosome duplication in GSCs or NBs induced to divide symmetrically. (A and B) Examples of *upd*-induced GSCs without *alms1*^*RNAi*^ (A, *nos-gal4*>*UAS-upd*) and with *alms1*^*RNAi*^ (B, *nos-gal4>UAS-upd, UAS-alms1^RNAi^*). Green: Vasa. Red: Asl. White: Alms1a. Blue: DAPI. Inset is magnified image of a single GSC. Arrows indicate centrosomes. Bar: 10 μm. (C) Quantification of centrosome numbers in the indicated genotypes. (D and E) Examples of NBs in *wor-gal4>UAS-apkc*^*CAAX*^ (D), *wor-gal4>UAS-apkc*^*CAAX*^, UAS-*alms1*^*RNAi*^ (E) brains. Green: Phalloidin. Red: Asl. White: Alms1a. Blue: DAPI. NBs are outlined with dotted lines. Arrowheads indicate NB centrosomes. Bar: 10 μm. (F) Quantification of centrosome numbers in the indicated genotypes. (G) Model of Alms1a’s requirement for daughter centriole duplication in stem cells.

We also examined whether symmetrically dividing NBs require *alms1a* for centriole duplication, by expressing the membrane-targeted aPKC (*aPKC*^*CAAX*^), which is known to induce symmetric NB divisions, leading to NB tumor formation (Lee et al., 2006). When *aPKC*^*CAAX*^ expression was combined with *alms1*^*RNAi*^, the NBs retained centrosomes despite efficient depletion of *alms1a* (Figures 7D-7F). Taken together, these results suggest that Alms1a is required for centriole duplication only in asymmetrically dividing cells.

## DISCUSSION

Asymmetric behaviors between mother and daughter centrosomes have been observed in asymmetrically dividing stem cells in several systems (Conduit and Raff, 2010; Habib et al., 2013; Januschke et al., 2011; Wang et al., 2009; Yamashita et al., 2007), provoking the idea that these centrosome asymmetries may contribute to asymmetric cell fates. Here, we identified a centrosomal protein Alms1a, a homolog of a causative gene of human ciliopathy Alstrome Syndrome, that exhibits asymmetric localization between mother and daughter centrosomes. Alms1a is required for centriole duplication specifically in asymmetrically dividing stem cells, but not in symmetrically dividing cells, demonstrating unique regulation of centrosomes in asymmetrically dividing stem cells. A recent study identified Ninein as a protein that specifically localizes to the daughter centrosome when overexpressed in *Drosophila* neuroblasts and male GSCs (Zheng et al., 2016), though consequences were not detected upon its depletion, leaving the role of its asymmetric localization unknown. Although the exact molecular mechanism of Alms1a function remains elusive, it is required for daughter centriole duplication only in asymmetrically dividing stem cells. A few lines of observations imply its possible function. 1) Alms1a localizes to the daughter centriole only at the mother centrosome of asymmetrically dividing stem cells. 2) *alms1a* depletion leads to failure in daughter centriole duplication, while the mother centriole gradually elongates. These results imply that Alms1a is required to ‘seed’ the production of daughter centriole. Considering that Alms1a is dispensable in symmetrically dividing cells, this Alms1a-dependent activity to ‘seed’ new centrioles is only required when two centrosomes (or two centrioles) are not equivalent (Figure 7G). Based on the fact the mother centriole in male GSCs elongates in the absence of *alms1a*, we further speculate that the mother centrioles in asymmetrically dividing stem cells might have a tendency to elongate, possibly due to their higher MTOC activity, necessitating a mechanism to counter such a tendency to allow the seeding of the new, daughter centrioles (Figure 7G).

In summary, our study demonstrates the essential role of *alms1a* for centriole duplication uniquely in asymmetrically dividing stem cells. This specific requirement of *alms1a* only in a subset of cell types (i.e. asymmetrically dividing adult stem cells) may partly explain the reported late-onset nature of Alstrom syndrome, which exhibits symptoms progressively during childhood, as opposed to many other ciliopathies that are associated with many symptoms at birth (Braun and Hildebrandt, 2017).

## Supporting information

Supplementary table-mass spectrometry

## ACKNOWLEDGEMENTS

We thank Drs. Cheng-Yu Lee, Jordan Raff, Cayetano Gonzales, the Bloomington Stock Center, the Vienna Drosophila RNAi Center, and the Developmental Studies Hybridoma Bank for reagents, and the Yamashita laboratory, Sue Hammoud and Life Science Editors for discussions and comments on the manuscript. MS Bioworks for mass-spectrometry analysis. This work was supported by R01GM118308 (to Y.M.Y). The research in the Yamashita laboratory is supported by Howard Hughes Medical Institute.

## AUTHOR CONTRIBUTIONS

CC, Conception and design, Acquisition of data, Analysis and interpretation of data, Drafting and revising the article. YMY, Conception and design, Supervision, Analysis and interpretation of data, Drafting and revising the article.

## DECLARATION OF INTERESTS

The authors declare no competing interests.

---

**Figure S1.**
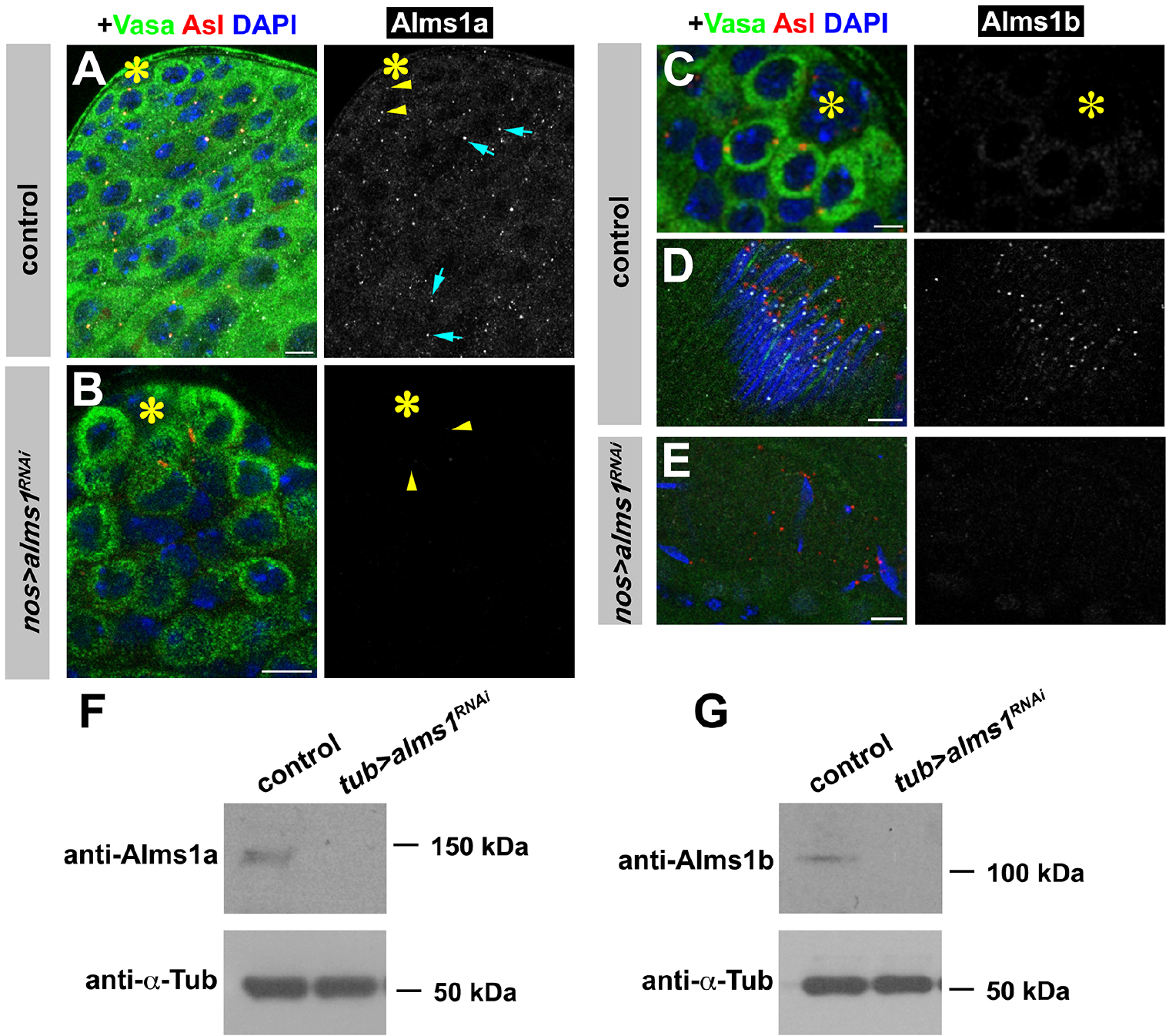
Validation of RNAi-mediated knockdown of *alms1* and antibody specificity for Alms1a and Alms1b. (A and B) Examples of Alms1a staining in control (A) and *nos-gal4>UAS-alms1*^*RNAi*^ (B) GSCs. Green: Vasa. Red: Asl. White: Alms1a. Blue: DAPI. Asterisk indicates the hub. Arrowheads (yellow) indicate GSC centrosomes. Arrows (cyan) indicate SG centrosomes. Bar: 10 μm. (C-E) Examples of Alms1b staining in control (C) GSCs, control (D) spermatids and *nos-gal4>UAS-alms1*^*RNAi*^ (E) spermatids. Green: Vasa. Red: Asl. White: Alms1a. Blue: DAPI. Bar: 5 μm. (F and G) Western blot analyses of Alms1a (F) or Alms1b (G) protein in the lysate of either control or *tub-gal4>UAS-alms1*^*RNAi*^ testes. α-Tubulin expression was used as a loading control.

**Figure S2.**
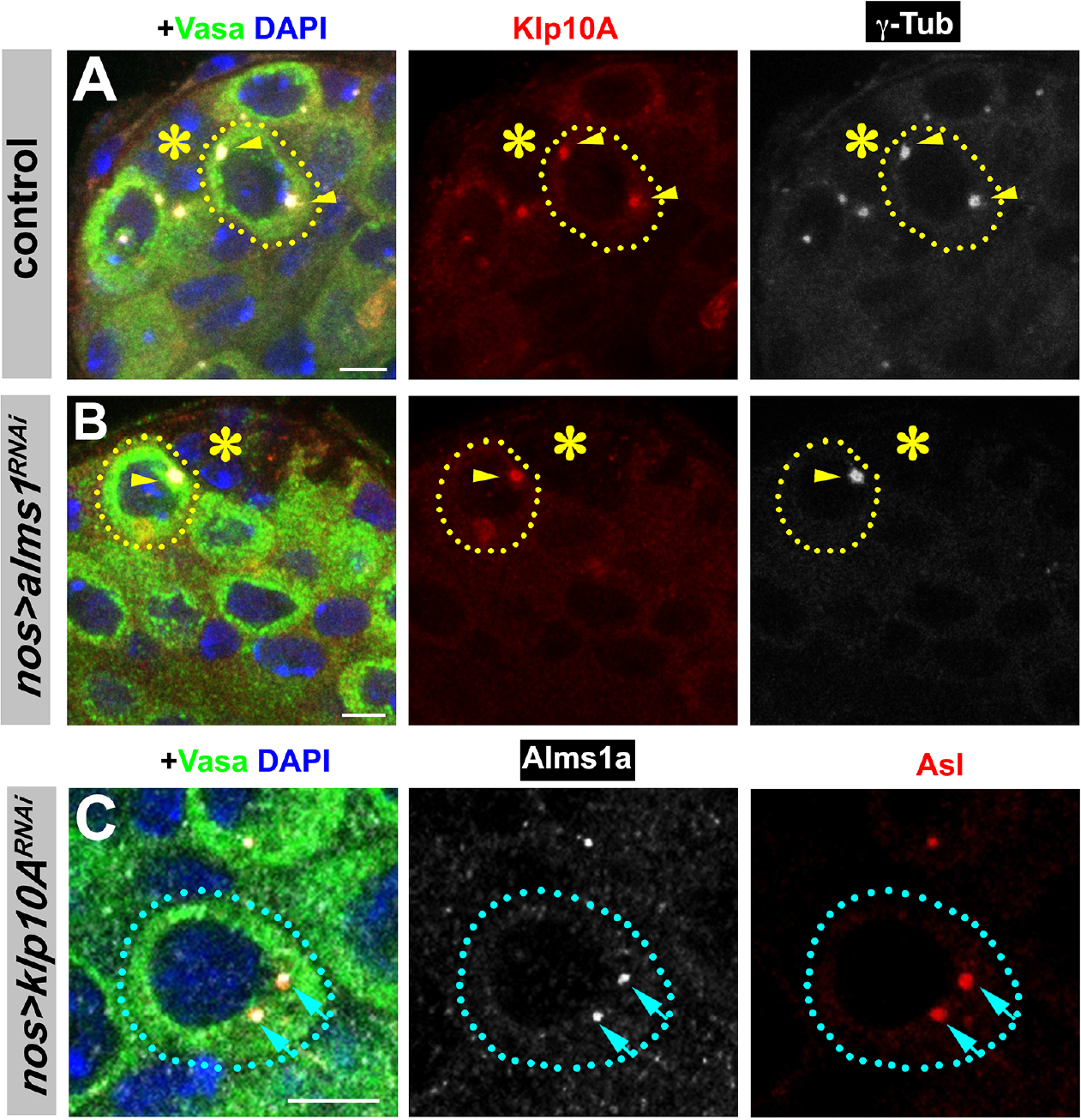
*alms1a* is not required for Klp10A localization to GSC centrosomes. (A and B) Localization of Klp10A in control (A) and *alms1*^*RNAi*^ (B) GSCs. Green: Vasa. Red: Klp10A. White: γ-Tub. Blue: DAPI. Note that *alms1*^*RNAi*^ results in centrosome loss, thus Klp10A localization was assessed on the remaining mother centrosome. Asterisk indicates the hub. Bar: 5 μm. (C) Localization of Alms1a remains intact in *klp10A*^*RNAi*^ SG. Green: Vasa. Red: Asl. White: Alms1a. Blue: DAPI. Asterisk indicates the hub. SG is indicated by a dotted line. Arrows indicate SG centrosomes. Bar: 5 μm.

**Figure S3.**
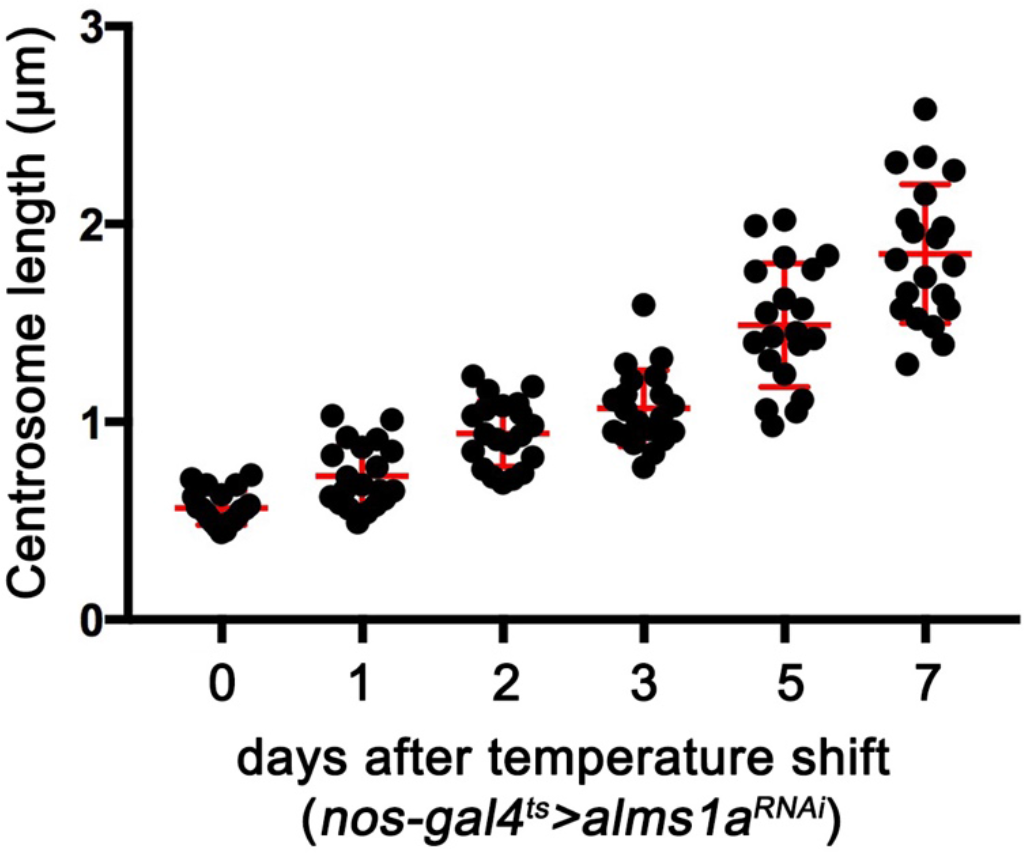
Quantification of GSC mother centrosome length after the induction of *alms1*^*RNAi*^. *nos-gal4, tub-gal80*^*ts*^>*UAS-alms1*^*RNAi*^ GSCs were shifted to 29°C to induce RNAi. Centrosome size was examined at 0-7 d after RNAi induction. Error bars indicate the standard deviation.

**Figure S4.**
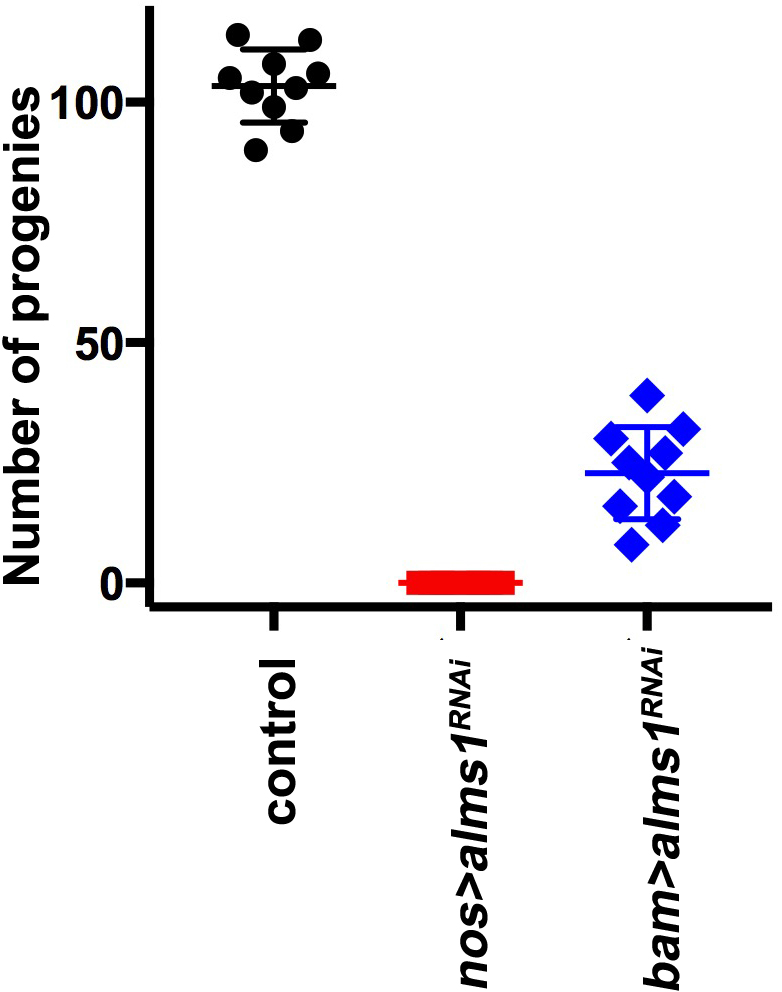
Fertility assays for control,*nos-gal4>UAS-alms1*^*RNAi*^ and *bam-gal4>UAS-alms1*^*RNAi*^. Error bars indicate the standard deviation. 10 independent crosses were used for each genotype. Note that the sterility of *nos-gal4>UAS-alms1*^*RNAi*^ flies likely reflects the depletion of both *alms1a* and *alms1b*. However, the depletion of *alms1* (*bam-gal4>UAS-alms1*^*RNAi*^) leads to milder fertility defect, suggesting that centrosome depletion in early germ cells due to *alms1a* depletion compromises fertility.

## Materials and Methods

### Fly husbandry, strains and transgenic flies

All fly stocks were raised on standard Bloomington medium at 25°C, and young flies (0− to 1− day-old adults) were used for all experiments unless otherwise noted. The following fly stocks were used: *nos-gal4* (Van Doren et al., 1998), *UAS-upd* (Zeidler et al., 1999), *UAS-dpp* (BDSC1486), *tub-gal80*^*ts*^ (McGuire et al., 2003), *UAS-klp10A-GFP* (Inaba et al., 2015), *UAS-GFP-α-tubulin84B* (Grieder et al., 2000), *UAS-klp10A*^*RNAi*^ (TRiP.HMS00920, BDSC), *UAS-alms1*^*RNAi*^ (TRiP.HMJ30289, BDSC), *nos-gal4* without VP16 (*nos-gal4ΔVP16*) (Inaba et al., 2015), *cnb-YFP* (Januschke et al., 2011), *ubi-asl-tdTomato* (Gopalakrishnan et al., 2011), *UAS-EGFP* (BDSC5430). Note that *UAS-alms1*^*RNAi*^ is expected to knockdown both *alms1a* and *alms1b*. However, because *alms1b* is not expressed in GSCs, the phenotypes resulting from expression of this RNAi construct in GSCs are due to the loss of *alms1a*. Although we attempted to generate *alms1a*-specific RNAi lines, none of them efficiently knocked down *alms1a* as evidenced by western blotting.

### Immunofluorescent staining and confocal microscopy

*Drosophila* testes were dissected in phosphate-buffered saline (PBS), transferred to 4% formaldehyde in PBS and fixed for 30 minutes. The testes were then washed in PBST (PBS containing 0.1% Triton X-100) for at least 30 minutes, followed by incubation with primary antibody in 3% bovine serum albumin (BSA) in PBST at 4°C overnight. Samples were washed for 60 minutes (three 20-minute washes) in PBST, incubated with secondary antibody in 3% BSA in PBST at 4°C overnight, washed as above, and mounted in VECTASHIELD with 4’,6-diamidino-2-phenylindole (DAPI; Vector Labs).

The primary antibodies used were as follows: rat anti-Vasa (1:20; developed by A. Spradling and D. Williams, obtained from Developmental Studies Hybridoma Bank (DSHB)); mouse anti-γ-Tubulin (GTU-88; 1:100; Sigma-Aldrich); rabbit anti-Vasa (d-26; 1:200; Santa Cruz Biotechnology); rabbit anti-Asl (1:2000; a gift from Jordan Raff); chicken anti-GFP (1:1000, Aves Labs). Anti-Alms1a and Alms1b antibody were generated by injecting peptides (QEMEVEPKKQLEKEQHQNDMQQGEPKGREC) and (CNISQRGNHLEKIE) into guinea pigs (Covance). Affinity purification was used to purify the antibodies. Specificity of the antibodies was validated by the lack of staining in *alms1*^*RNAi*^ testis (Figure S1). Anti-Centrobin antibody was generated by injecting a peptide (GRPSRELHGMVHSTPKSGSVEPLRHRPLDDNIC) into rabbits (Covance). Alexa Fluor-conjugated secondary antibodies (Life Technologies) were used with a dilution of 1:200. Images were taken using an upright Leica TCS SP8 confocal microscope with a 63× oil immersion objective (NA=1.4) and processed using Adobe Photoshop software. Superresolution imaging was performed on Leica TCS SP8 with Hybrid spectral dectectors and Hyvolution.

### Co-immunoprecipitation and western blotting

For immunoprecipitation using *Drosophila* testis lysate, testes enriched with GSCs due to ectopic Upd expression (100 pairs/sample) were dissected into PBS at room temperature within 30 min. Testes were then homogenized and solubilized with lysis buffer (10 mM Tris-HCl pH 7.5; 150 mM NaCl; 0.5 mM EDTA supplemented with 0.5% NP40 and protease inhibitor cocktail (EDTA-free, Roche)) for 30 min at 4°C. Testes lysates were centrifuged at 13,000 rpm for 15 min at 4°C using a table centrifuge, and the supernatants were incubated with GFP-Trap magnetic agarose beads (ChromoTek) for 4 h at 4°C. The beads were washed three times with wash buffer (10 mM Tris-HCl pH 7.5; 150 mM NaCl; 0.5 mM EDTA). Bound proteins were resolved in SDS-PAGE and analyzed by western blotting using anti-Alms1a and anti-GFP antibodies.

For anti-Klp10A pull-down and mass spectrometry assays, testes enriched with GSCs (*nos-gal4>UAS-upd*) or enriched with SGs (*nos-gal4>UAS-dpp*) (250 pairs/sample) were dissected into PBS. Testes were then homogenized and solubilized with lysis buffer (10 mM Tris-HCl pH 7.5; 150 mM NaCl; 0.5 mM EDTA, 0.1% NP40 and protease inhibitor cocktail (EDTA-free, Roche)) for 30 min at 4°C. After centrifugation at 13,000 rpm for 15 min at 4°C, supernatants were incubated with anti-Klp10A conjugated Dynabeads for 2h at 4°C. The beads were washed three times with wash buffer (10 mM Tris-HCl pH 7.5; 150 mM NaCl; 0.5 mM EDTA), followed by boiling in 2xLaemmli buffer (Bio-Rad) and sent for Mass spectrometry analysis (MS Bioworks).

For western blotting, samples subjected to SDS-PAGE (NuPAGE Bis-Tris gels (8%; Invitrogen)) were transferred onto polyvinylidene fluoride (PVDF) membranes (Immobilon-P; Millipore). Membranes were blocked in PBS containing 5% nonfat milk and 0.1% Tween-20, followed by incubation with primary antibodies diluted in PBS containing 5% nonfat milk and 0.1% Tween-20. Membranes were washed with PBS containing 5% nonfat milk and 0.1% Tween-20, followed by incubation with secondary antibody. After washing with PBS, detection was performed using an enhanced chemiluminescence system (Amersham). Primary antibodies used were rabbit anti-GFP (abcam; 1:4000), guinea pig anti-Alms1a tag (this study; 1:5000). Secondary antibodies used were goat anti-guinea pig IgG and goat anti-rabbit IgG conjugated with horseradish peroxidase (HRP) (abcam; 1:5000).

### Data analyses

Statistical analysis was performed using GraphPad Prism 7 software. Data are shown as means±standard deviation. The *P*-value (two-tailed Student’s *t-test*) is provided for comparison with the control. Mother and daughter centriole size analysis was performed by manually drawing regions of interest in ImageJ.

### Fertility assays

Individual males of the genotypes shown were allowed to mate with virgin female *yw* flies for 1 week. The flies were removed, and the numbers of progenies were scored.

